# Genetic basis of R-phase H:z_27_ antigen expression and phylogenetic relationships of *Salmonella enterica* serovar Senftenberg

**DOI:** 10.1101/2025.10.27.684867

**Authors:** Chieko Kato, Takahiro Ochiai, Yuta Ohno, Ikuo Uchida, Nobuo Arai, Hidemasa Izumiya, Tetsuya Ikeda, Masahiro Kusumoto, Masato Akiba

## Abstract

*Salmonella enterica* serovar Senftenberg (*S*. Senftenberg) is one of the most prevalent *Salmonella* serovars and can be isolated from various animals and humans. According to the White–Kauffmann–Le Minor (WKL) scheme, the antigenic formula of *S*. Senftenberg is defined as 1,3,19:g,[s],t:–. Although the antigenic formula 1,3,19:z_27_:– was formerly designated as serovar Simsbury, this name was later removed from the WKL scheme due to reports of interconvertibility between serovars Senftenberg and Simsbury. The H:z_27_ antigen is currently recognized as one of the R-phase H antigens of *S*. Senftenberg. To elucidate the genetic basis of H:z_27_ antigen expression in *S*. Senftenberg, we determined the whole-genome sequences of isolates expressing the H:z_27_ antigen. These isolates belonged to sequence type (ST) 185 and harbored a horizontally acquired chromosomal region containing the *fljA*, *fljB*, and *pinR* genes, which are predicted to encode a translational repressor of *fliC* mRNA, a phase 2 flagellin, and an invertase, respectively. A deletion mutant lacking this region expressed the H:g,s,t antigen instead of H:z_27_, as confirmed by serotyping and Western blot analysis. Phylogenetic analysis based on *S*. Senftenberg genome archives, together with the sequences obtained in this study, suggested that the *S*. Senftenberg ST185 lineage acquired the *fljAB*-*pinR* locus around 1867. In contrast, our broader phylogenetic analysis of *Salmonella* serovars indicated that *S*. Senftenberg ST14 is genetically closer to certain other serovars than to ST185, while ST185 shows a closer relationship to a different serovar than to ST14. These findings suggest that *S*. Senftenberg comprises at least two distinct genomic lineages.

**IMPORTANCE:** Limited information is available on the genetic basis of R-phase H antigen expression in *S. enterica*. In this study, we first demonstrated that the R-phase H:z_27_ antigen of *S.* Senftenberg is determined by a horizontally acquired chromosomal region. The ST 185 lineage of *S.* Senftenberg appears to have acquired this chromosomal region around 1867 and subsequently disseminated worldwide. We also demonstrated that *S.* Senftenberg comprises at least two distinct genomic lineages. The combined use of serotyping and multilocus sequence typing may provide a typing framework that better reflects the disease potential of *Salmonella*.

## INTRODUCTION

The genus *Salmonella* comprises Gram-negative rods belonging to the family Enterobacteriaceae and includes two species: *S. enterica* and *S. bongori*. *S. enterica* is further divided into six subspecies: *enterica*, *salamae*, *arizonae*, *diarizonae*, *houtenae*, and *indica*. Beyond this taxonomic classification, serotyping is widely used as an epidemiological typing method to further subdivide *Salmonella* species. In the White–Kauffmann–Le Minor (WKL) scheme, serovars are defined based on combinations of cell surface lipopolysaccharide components (O antigens) and one or two distinct flagellar proteins (H antigens) (1). To date, more than 2,600 serovars have been identified within the genus *Salmonella* (2, 3).

To enable finer discrimination of individual serovars, molecular subtyping methods have been developed, including pulsed-field gel electrophoresis, multiple-locus variable-number tandem repeat analysis, and multilocus sequence typing (MLST) (4). In recent years, the accumulation of whole-genome sequence data in public databases has led to the growing use of MLST-based genotyping (5). Concurrently, there is increasing interest in revisiting conventional serotyping approaches in light of whole-genome sequence information (6).

*S. enterica* subsp. *enterica* is a major cause of gastroenteritis in both humans and animals. *S. enterica* serovar Senftenberg (*S*. Senftenberg) is one of the most prevalent and can be isolated from various food sources, including poultry, beef, and seafood worldwide (7). In the United States, this serovar ranks among the top five serovars isolated from food and the top eleven serovars isolated from clinically ill animals. Human infections are typically associated with exposure to farm environments or the consumption of contaminated food (8). Strains of *S*. Senftenberg are primarily classified into two major multilocus sequence types (STs): ST14 and ST185 (9). These STs share no common alleles across the seven loci used in MLST typing, suggesting a substantial genetic distance between ST14 and ST185 (8).

The antigenic formula of *S.* Senftenberg is defined as 1,3,19:g,[s],t:–. According to the WKL scheme, this serovar may also express R-phase H antigens instead of g,[s],t, including z_27_, z_34_, z_43_, z_45_, or z_46_ (1). The “R phase” of H antigen was first described by Kauffmann and refers to abnormal specificities of H antigens (10). Although the antigenic formula 1,3,19:z_27_:– was formerly designated as serovar Simsbury, the name was later removed from the scheme due to reports of interconvertibility between serovars Senftenberg and Simsbury (11). However, the molecular mechanisms underlying the expression of the H:z_27_ antigen and the STs to which the strains belong remain unclear.

This study was conducted to elucidate the genetic basis of H:z_27_ antigen expression and to investigate the phylogenetic relationships among *S.* Senftenberg strains expressing the typical or R-phase H antigens.

## MATERIALS AND METHODS

### Bacterial isolation and identification

The *S.* Senftenberg isolates used in this study are listed in Table S1. A total of 18 isolates were collected in Japan for diagnostic or monitoring purposes by staff at local animal hygiene service centers, public health institutes, or the Food and Agricultural Materials Inspection Center. One exception is isolate SL212, which had been stored at the National Institute of Infectious Diseases, Japan, and whose origin remains unknown. Patient information was anonymized and de-identified prior to analysis.

The isolates were epidemiologically unrelated and were obtained from distinct samples collected in 7 of the 47 prefectures across Japan. *Salmonella* spp. were identified based on colony morphology on selective media and biochemical characteristics, as previously described (12). Serovar identification was conducted using microtiter and slide agglutination methods according to the WKL scheme (1). All isolates were stored at –80 °C until further analysis.

### Growth conditions and genomic DNA extraction

All bacterial strains were cultured in Tryptic Soy Yeast Extract Broth at 37 °C under static conditions for 20-24h. Genomic DNA was extracted from bacterial cultures using the DNeasy Blood & Tissue Kit (QIAGEN, Hilden, Germany), according to the manufacturer’s instructions. For elution, Buffer EB was used instead of Buffer AE. DNA concentration was measured using a Qubit 2.0 Fluorometer (Thermo Fisher Scientific, Waltham, MA).

### Short-read sequencing

Short-read sequencing was conducted on the Illumina MiSeq platform (Illumina, San Diego, CA). Genomic DNA libraries were constructed using the QIAseq FX DNA Library Kit (QIAGEN), according to the manufacturer’s instructions. Sequencing was carried out with the MiSeq Reagent Kit v3 (600 cycles), producing 300 bp paired-end reads. Raw reads were quality-filtered and trimmed using fastp v0.23.4 (13) with default settings.

### In silico MLST

Draft genome sequences were *de novo* assembled from fastp-processed Illumina MiSeq reads using SKESA v2.4.0 (14). Subsequently, *in silico* MLST was performed using mlst v2.23.0 (https://github.com/tseemann/mlst).

### Long-read sequencing

Long-read sequencing was performed using the Oxford Nanopore MinION Mk1C platform (Oxford Nanopore Technologies, Oxford, UK). Genomic DNA libraries for Nanopore sequencing were prepared according to the manufacturer’s instructions. DNA repair and end-preparation were conducted using the NEBNext Companion Module for Oxford Nanopore Technologies Ligation Sequencing (New England Biolabs, Ipswich, MA). Barcoding and adapter ligation were subsequently carried out using the Ligation Sequencing Kit (Oxford Nanopore Technologies) and the Native Barcoding Expansion 1–12 (Oxford Nanopore Technologies). Magnetic bead-based purification was performed using AMPure XP beads (Beckman Coulter, Brea, CA). Final wash steps were conducted using LFB buffer (Oxford Nanopore Technologies) as an alternative to ethanol. The prepared libraries were then loaded onto an R9.4.1 flow cell (Oxford Nanopore Technologies) and sequenced using the MinION device. After sequencing, raw Nanopore reads were quality-filtered and trimmed using NanoFilt v2.8.0 (15). For each strain, reads were processed to ensure high quality: reads with a quality score below 10 or a length shorter than 500 bp were discarded, and the first 75 bases of each read were trimmed to remove potential adapter sequences.

### Hybrid genome assembly

Hybrid genome assembly was performed using Unicycler v0.4.8 (16, 17) for strains. Quality-filtered short reads from the Illumina MiSeq platform and processed long reads from the Oxford Nanopore MinION were used as input for the assemblies. Genome annotation was performed using DFAST v1.6.0 (https://dfast.ddbj.nig.ac.jp/) for the purpose of public database submission, with taxonomic rank set to species and the taxon to *S. enterica*. In parallel, Prokka v1.14.6 (18) was used to identify flagellar genes, including *fliC*, *fljB*, and other associated loci, and to characterize their genomic context. In cases where the *fljB* gene was identified, a neighboring coding sequence annotated as a hypothetical protein was further examined. BLASTp searches were performed using the UniProtKB web interface (https://www.uniprot.org/blast), which revealed high similarity to the *fljA* gene product (translational repressor of phase-1 flagellin), supporting its functional assignment. Accordingly, these loci were manually annotated as *fljA* in the final genome annotation used for analysis.

### Construction of *fljAB-pinR* deletion mutant of *Salmonella* serovar Senftenberg

All primers used in this study were purchased from FASMAC (Kanagawa, Japan) and are listed in Table S2. The primer pair FA-F and FA-R was used to amplify the upstream region of the *pinR* gene (fragment A), and the primer pair FB-F and FB-R was used to amplify the downstream region of the *fljB* gene (fragment B) from *S.* Senftenberg strain HOLHSC-37-53 by PCR. The 5′ terminal 25 nucleotides of the FA-R primer were designed to be reverse complementary to the FB- F primer sequence. Overlap extension PCR was then performed using the primer pair FA-F and FB-R to join fragments A and B. The resulting fused fragment was digested with BamHI and SalI, cloned into the pTH18cs1 vector (19), and used for targeted deletion of the *fljB* gene via gene replacement.

*S*. Senftenberg strain HOLHSC-37-53 was transformed with the gene replacement vector by electroporation. The cells were spread on Luria–Bertani (LB) agar plates (Becton, Dickinson and Co., Franklin Lakes, NJ, USA) supplemented with chloramphenicol and incubated at 28 °C for 18 h. The resulting colonies were then streaked onto prewarmed LB agar plates and incubated at 42 °C for 18 h. Single-crossover strains were purified under the same conditions and subsequently passaged several times at 28 °C. The primer pair FljB-F and FljB-R was used to screen for double-crossover strains. Deletion of the *fljB* gene was confirmed by PCR amplification of the deleted region using the primer pair CU and CD, followed by sequencing of the amplified fragments.

### Preparation of flagella and western blot analysis

Bacterial cultures were grown at 37°C in LB broth with shaking at 250 rpm for 16 hours. The cultures were then centrifuged at 6,000 × g for 20 minutes, and the resulting supernatant was passed through a 0.22-μm pore size syringe filter. To recover sheared flagellar proteins, trichloroacetic acid (10% vol/vol) was added to the supernatant, and the mixture was incubated on ice for 30 minutes. The precipitated protein pellet was washed with ice-cold ethanol and resuspended in phosphate-buffered saline (PBS). Equal amounts of protein were loaded onto 12.5% SDS–polyacrylamide gels (cPAGEL HR; ATTO, Japan) and separated by electrophoresis using an AE-7300 electrophoresis system (ATTO).

Following SDS-PAGE, proteins were transferred onto PVDF membranes (Bio-Rad Laboratories, Hercules, CA) by semi-dry blotting using the WSE-4115 PoweredBlot Ace system (ATTO) with a standard transfer buffer containing 48 mM Tris, 39 mM glycine (pH 8.3), 0.0375% SDS, and 20% methanol. Membranes were blocked with 5% Block Ace (Yukijirushi Nyugyo, Tokyo, Japan) in TBS-T (Tris-buffered saline containing 0.1% Tween-20) for 1 h at room temperature. Subsequently, the membranes were incubated for 1 h with one of the following primary antibodies at appropriate dilutions: polyclonal rabbit anti-z_27_ antiserum (SSI Diagnostica, Hillerød, Denmark) or rabbit anti-FliC (H-G) antiserum (Denka, Tokyo, Japan). After washing, bound antibodies were detected using horseradish peroxidase (HRP)-conjugated anti-rabbit IgG antibodies and visualized with the Enhanced Chemiluminescence Prime Western Blotting System (GE Healthcare Technologies, Chicago, IL).

### Core Genome SNP Detection and Phylogenetic Analysis

Phylogenetic analysis of *S*. Senftenberg isolates was performed using the method described below, utilizing a total of 268 reference genome assemblies, as listed in Tables S1 and S3. Short reads were mapped to the complete genome of *S.* Senftenberg strain FSW0104 (accession no. CP037894.1) using Snippy v4.6.0 (https://github.com/tseemann/snippy), followed by construction of a core genome alignment and SNP calling. Intact prophage regions in the reference genome were identified with PHASTER (20) and masked prior to analysis. Recombination regions were detected and removed using Gubbins v3.1.6 (21), and core genome SNPs (cgSNPs) were extracted with SNP-sites v2.5.1 (22). A maximum-likelihood (ML) phylogenetic tree was generated from the concatenated cgSNP alignment using RAxML-NG v1.1 (23) with 1,000 bootstrap replicates, and visualized in iTOL v6 (24).

Genes associated with phase variation were identified by performing nucleotide BLAST (blastn) v2.16.0 (25) with the following thresholds: minimum length coverage >90% and nucleotide sequence identity >90%. The query sequences used were *fljA*, *fljB*, and *hin* (accession number AP044013).

Phylogenetic analysis of *S*. enterica strains was conducted using the method described below, incorporating 38 reference genome assemblies listed in Table S4, together with the genome sequence data of *S*. Senftenberg strains HOLHSC-37-53 and L-1930 obtained in this study. Whole-genome sequences were analyzed using CSI Phylogeny v1.4 (https://cge.food.dtu.dk/services/CSIPhylogeny/) with default parameters, which includes SNP calling, filtering, and construction of a core genome alignment. A ML phylogenetic tree was generated based on the concatenated SNP alignment, and visualized using iTOL v6.

### Temporal Bayesian phylogenetic analysis

To perform a temporal analysis of 87 *S*. Senftenberg strains, cgSNPs were extracted as described above. The transversion model of nucleotide substitution was selected as the best-fitting model according to the Bayesian Information Criterion using jModelTest 2.1.10 (26, 27). We evaluated twelve combinations of six clock models and two types of tree priors. The isolation year of each strain was used to calibrate the time scale of the phylogenetic tree. For strain SL212, the analysis was performed under the assumption that the isolation year was 1950, as the exact year is not known. Markov chain Monte Carlo analyses for each parameter combination were performed with a chain length of 10^8^. The best-fitting parameter combination was selected based on the marginal likelihood estimates obtained via the Path Sampler tool packaged in BEAST 2.6.7 (28), resulting in the combination of the “optimized relaxed clock” (clock model) and “coalescent: exponential population” (tree prior). Triplicate BEAST runs were conducted, and convergence was confirmed by ensuring all effective sample size values exceeded 200 using Tracer v1.7.2 (29). The resulting three tree files were summarized using LogCombiner v2.6.7, and a single maximum clade credibility tree was generated with TreeAnnotator v2.6.7, followed by visualization with FigTree v1.4.4 (https://github.com/rambaut/figtree/).

## RESULTS

### Characteristics of *S*. Senftenberg isolates expressing H:z_27_ antigen

We isolated four *S*. Senftenberg isolates expressing the H:z_27_ antigen from four distinct cases of bovine salmonellosis in Japan between 2021 and 2023. All isolates belonged to ST185. The H:z_27_ antigen was detected in each isolate; however, induction of the alternate phase of the H antigen was unsuccessful using the standard serotyping method. Consequently, the antigenic formula of these isolates was determined to be O1,3,19:z_27_:–.

### Identification of flagellar antigen coding regions

In this study, we determined the complete genome sequences of four *S.* Senftenberg isolates: two ST185 isolates expressing H:z_27_ antigens and two ST14 isolates expressing H:g,s,t antigens. Comparative analysis of these genomes with available *S*. Senftenberg genome sequence archives allowed us to identify chromosomal regions responsible for H antigen expression. As shown in Figure 1, the *fliC* gene, which encodes the phase 1 flagellar antigen, and its flanking regions were identical among the isolates (Fig. 1A). In contrast, the ST185 strains HOLHSC-37- 53 and HKLHSC-R3-12, possessed an additional chromosomal region predicted to regulate expression of a second phase H antigen. This region comprises three genes: *fljA*, *fljB*, and *pinR*, which are predicted to encode a translational repressor of *fliC* mRNA, a phase 2 flagellin, and an invertase, respectively. These genes are present in a subset of *S*. Senftenberg ST185 isolates, including FSW0104 (Fig. 1B). With the exception of *S*. Senftenberg ST185 strains, *Escherichia coli* strain ECC-1470, isolated from bovine mastitis in the USA, was the only strain found to possess a sequence exhibiting 100% identity to this region with complete coverage in the database. No additional genomic regions potentially associated with alternative H antigen expression were identified in the genome sequences of the *S*. Senftenberg strains.

**Figure 1.**
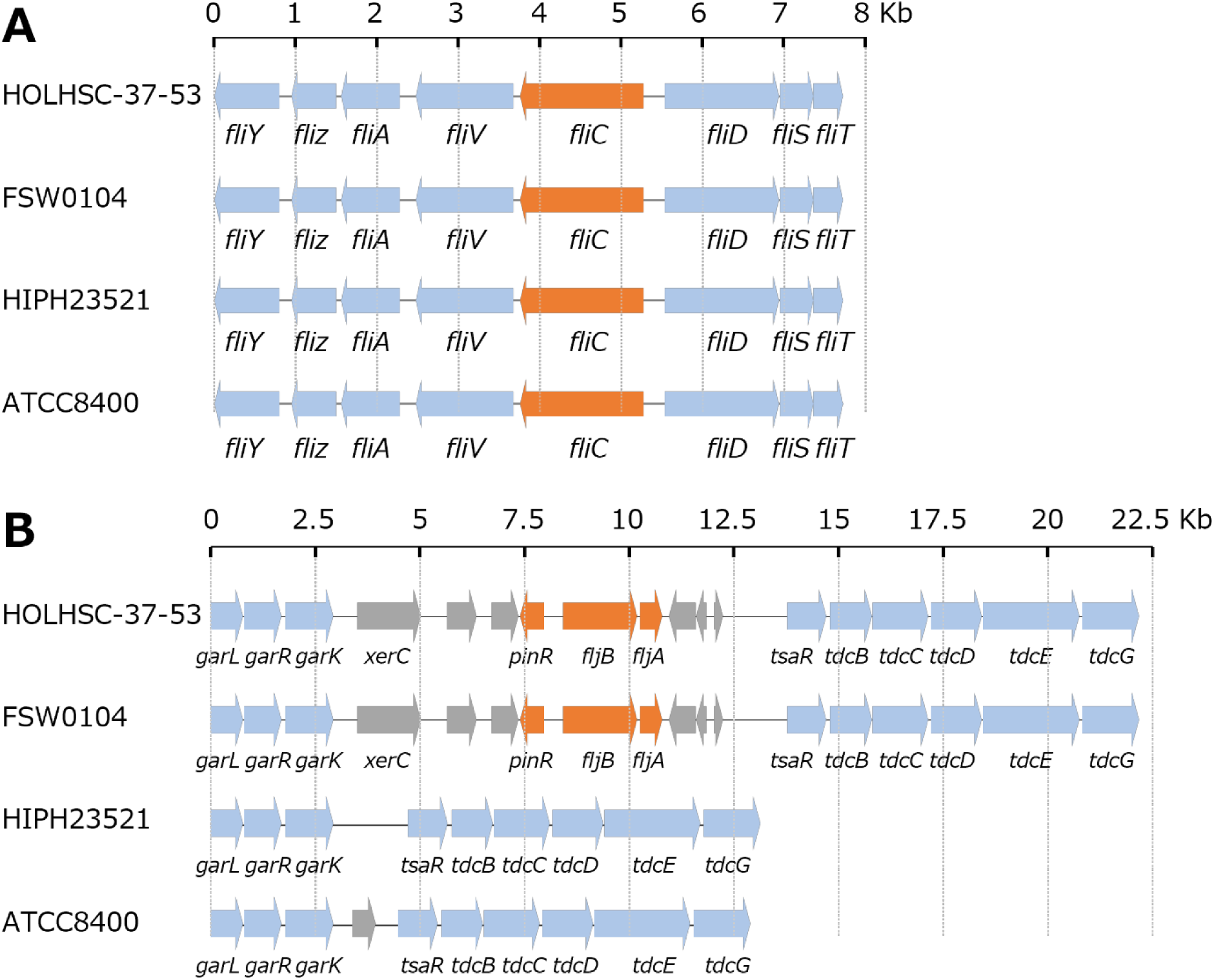
Schematic diagrams of chromosomal regions of *S. enterica* serovar Senftenberg isolates associated with phases 1 (A) and phase 2 (B) flagellar expression. The STs of isolates HOLHSC-37-53, FSW0104, HIPH23521, and ATCC8400 are ST185, ST185, ST14, and ST290, respectively. Orange arrows indicate genes involved in flagellin expression, whereas gray arrows indicate genes absent from two or more of the four isolates.

### Characteristics of *fljAB*-*pinR* deletion mutant

A *fljAB*-*pinR* deletion mutant of *S.* Senftenberg HOLHSC-37-53 was successfully constructed. The mutant strain retained motility, and its H antigen reacted with g,s,t antiserum but not with z_27_ antiserum.

### Western blot analysis of *fljAB-pinR* deletion mutant

SDS-PAGE analysis of the flagellar protein preparation from *S*. Senftenberg HOLHSC- 37-53 revealed a dominant protein band at approximately 61 kDa and a weaker band at 52 kDa. In contrast, the *fljAB*-*pinR* deletion mutant of the same strain exhibited a dominant band at 52 kDa and lacked the 61 kDa band (Fig. 2A). Western blot analysis showed that the 61 kDa band from the wild-type strain reacted specifically with anti-z_27_ serum (Fig. 2B), whereas both the 61 kDa and 52 kDa bands reacted with H-G serum (Fig. 2C). Given that the predicted molecular masses of FljB and FliC, based on their amino acid sequences, are 61.3 kDa and 51.6 kDa, respectively, these results indicate that the z_27_ antigen is encoded by the *fljB* gene.

**Figure 2.**
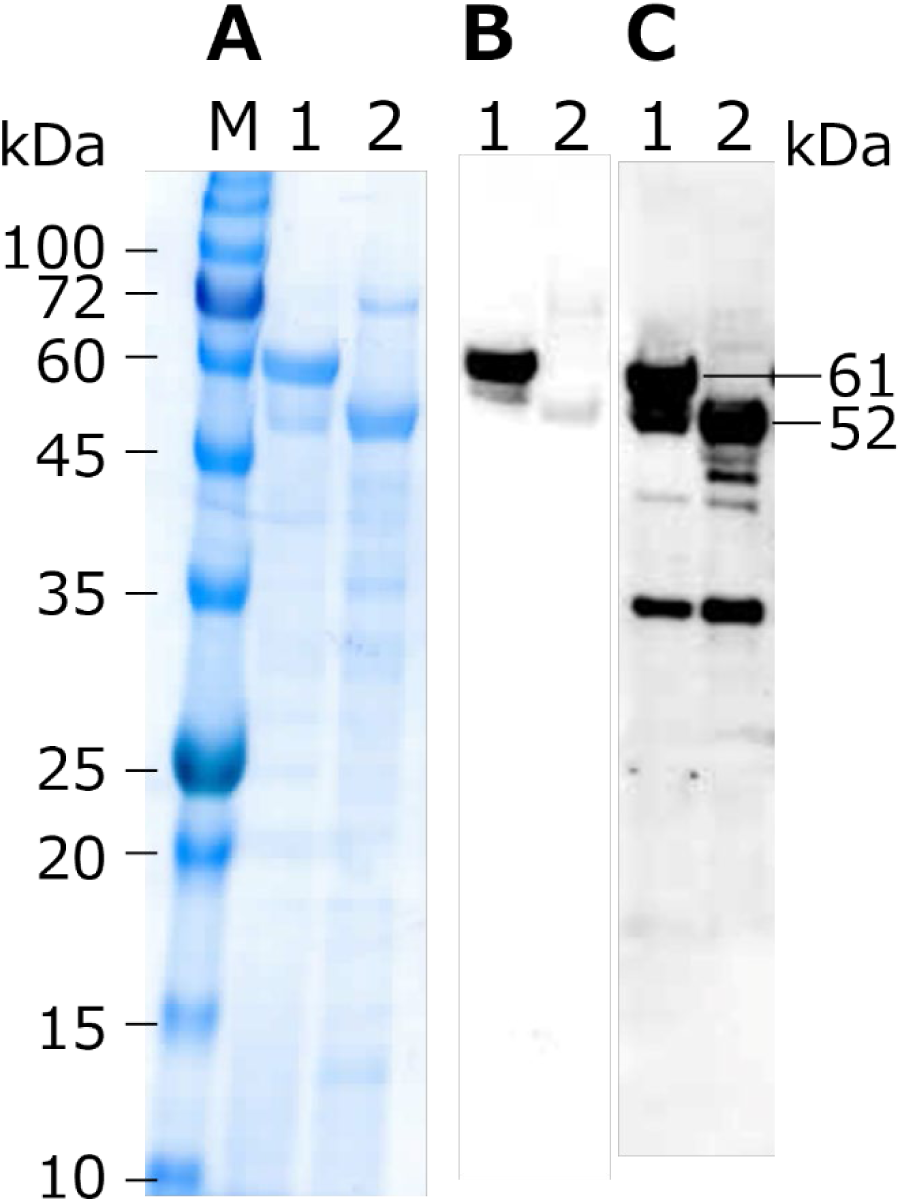
SDS-PAGE and western blot images demonstrating the lack of H:z_27_ antigen expression of the mutant strain. Proteins present in culture supernatants were stained by CBB (A). Anti-H:z_27_ serum (B) and anti-H:G serum (C) were used to detect proteins transferred to the PVD membrane. Lane M, protein ladder marker; Lane 1, wild-type strain, HOLHSC-37-53; Lane 2, *fljAB*-*pinR* deletion mutant of HOLHSC-37-53.

### Phylogenetic analysis of *S.* Senftenberg isolates

Figure 3 shows a circular phylogenetic tree constructed by using 250 *S*. Senftenberg genome sequence archives in addition to those of the 18 *S*. Senftenberg Japanese isolates analyzed by ourselves. Strains of ST14 and ST185 accounted for 73% of the *S* Senftenberg strains used in this study. The two STs formed distantly related clades in the phylogenetic tree. Strains of ST217 formed a clade with ST185, whereas strains of ST 210 formed a different clade with ST 14. The *fljAB*-*pinR* region distributed among 44 out of 69 ST185 strains. Five strains containing *fljB* like sequences with more than 95% identity were distributed in the clades of ST14, ST185, and ST210.

**Figure 3.**
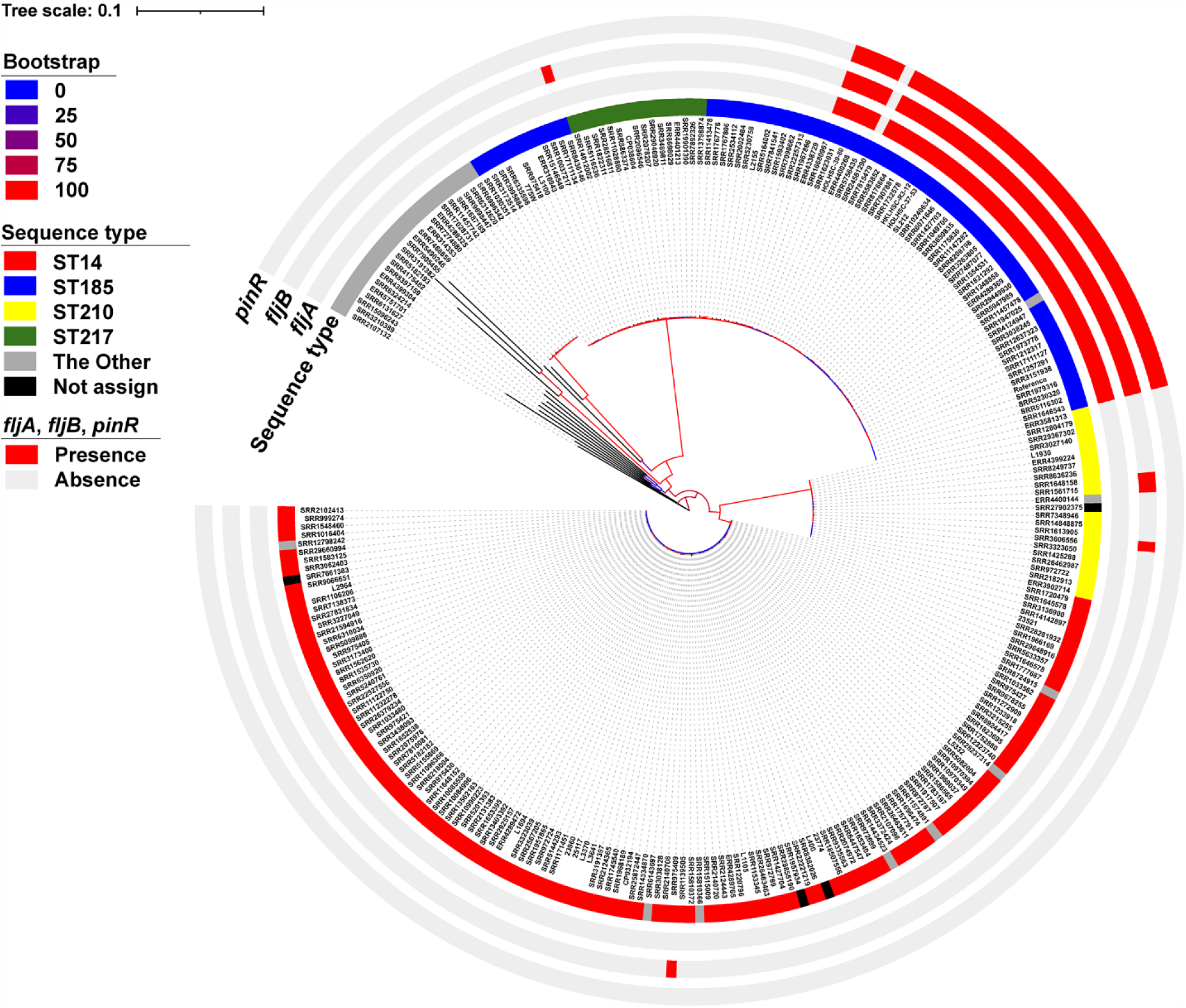
Phylogeny of *S. enterica* serovar Senftenberg isolates representing different sequence types (STs), including those expressing the H:z_27_ antigen. A phylogenetic tree was reconstructed based on 45,085 core-genome SNPs identified among 267 *S*. Senftenberg strains and the reference strain FSW0104. From the inner to the outer rings, the colored rings indicate STs (based on MLST) and the presence of *fljA*, *fljB*, and *pinR*.

Temporal Bayesian phylogenetic analysis of *S.* Senftenberg isolates belonging to ST185, ST217, and ST2653 revealed that the ST185 lineage harboring *fljAB*-*pinR* region diverged from the common ancestor around 1867 (95% Highest Posterior Density interval: 1816-1911) (Fig. 4).

**Figure 4.**
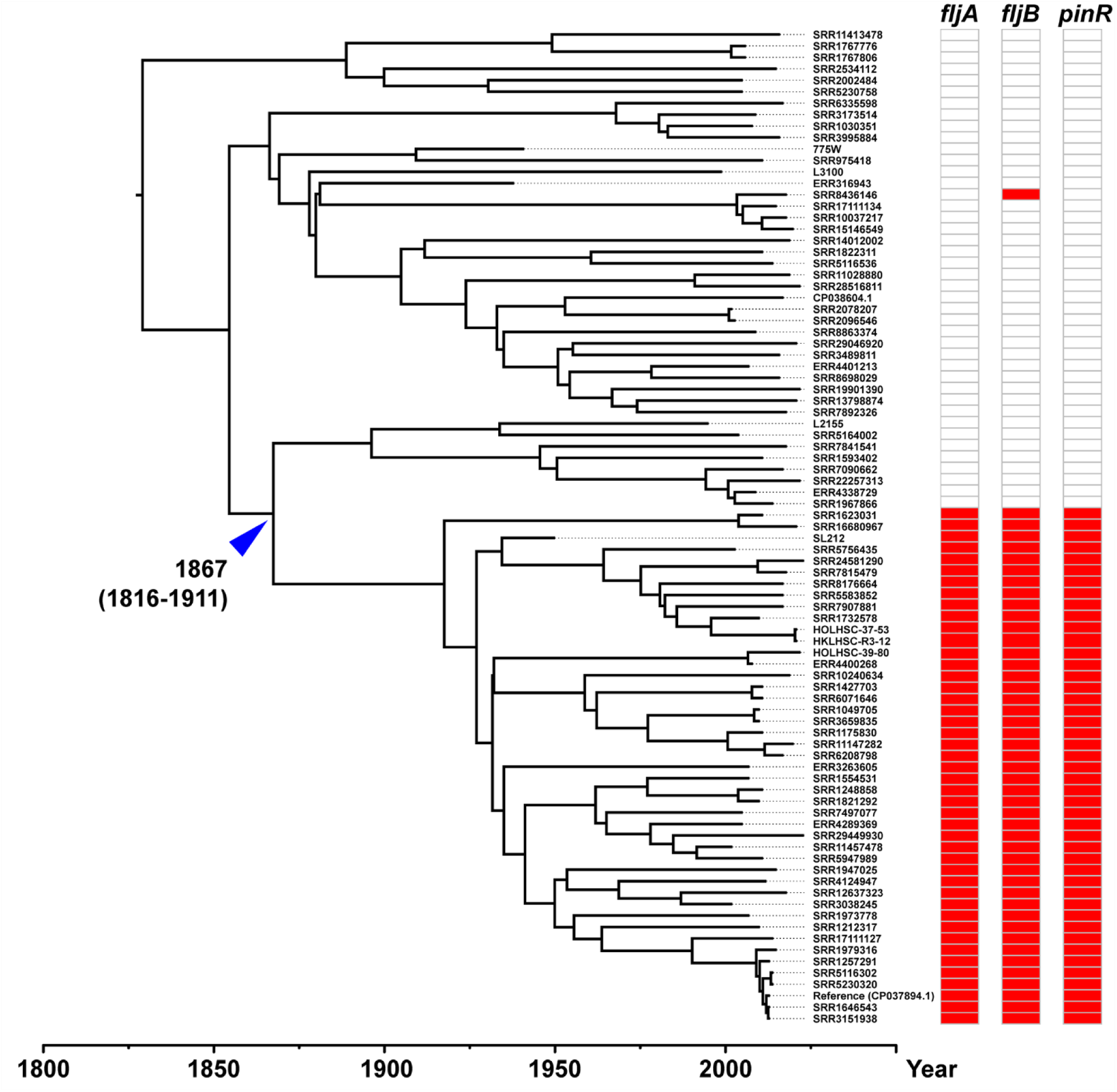
Temporal Bayesian phylogenetic analysis of *S. enterica* serovar Senftenberg isolates belonging to ST185, ST217, and ST2653. A phylogenetic tree was constructed using BEAST v2.6.7 based on 37,360 concatenated SNPs derived from the core-genome regions of 87 strains. The presence of *fljA*, *fljB*, and *pinR* is indicated on the right. The x-axis represents the estimated time of emergence of each strain. The predicted acquisition time of the *fljAB-pinR* locus is shown by a blue arrow, with the 95% highest posterior density interval indicated.

### Phylogenetic position of *S*. Senftenberg STs among *S. enterica* serovars

Figure 5 presents a phylogenetic tree constructed from genome sequence data of 40 *S. enterica* strains representing 27 different serovars, together with *S*. Senftenberg strains HOLHSC37-53 and L-1930 (ST210), whose genome sequences were determined in this study. Among the *S*. Senftenberg isolates, ST217 and ST290 clustered with ST185, whereas ST210 clustered with ST14. In contrast, a substantial genetic distance was observed between ST185 and ST14. For reference, the allele number profiles of *S*. Senftenberg STs are listed in Table S5. *S*. Senftenberg ST185 formed a clade with *S. enterica* serovars Dessau and Kentucky, whereas ST14 formed a clade with serovars Albany, Tennessee, Rissen, Goldcoast, and Agona.

**Figure 5.**
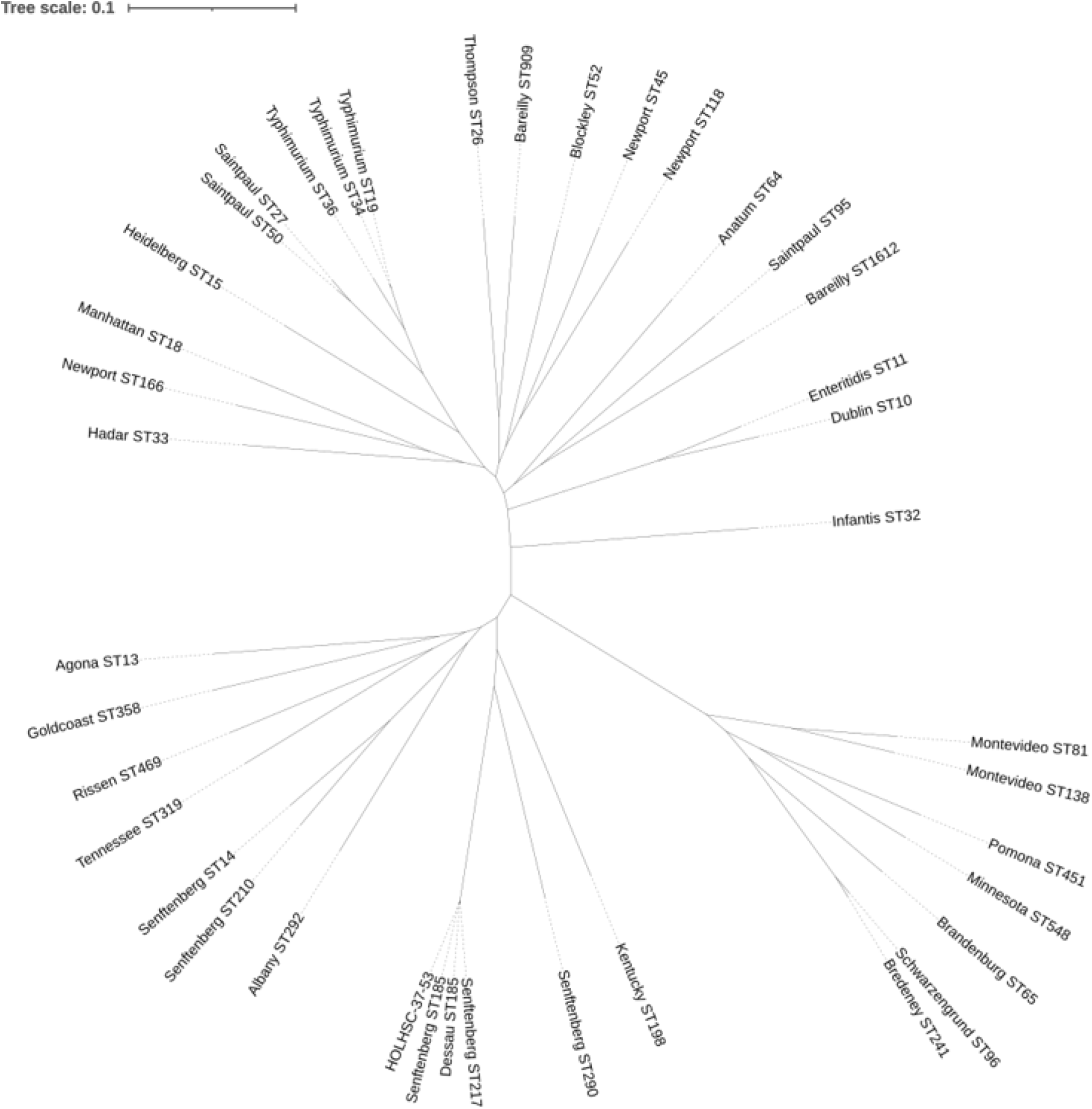
Phylogeny of *S. enterica* serovars. A phylogenetic tree was constructed using iTOL v6 based on 144,051 concatenated SNPs from the core-genome regions of 40 *S. enterica* strains.

## DISCUSSION

Many serovars of *S. enterica* have the ability to express two different H antigens, but only one type of antigen is expressed by a single cell at any given time (30). Phase 1 and phase 2 flagellins are encoded by the *fliC* and *fljB* genes, respectively, which are located at different loci on the chromosome. The *fljB* gene is located between the *fljA* and *hin* genes, and the switch in flagellin expression between phases 1 and 2 is regulated by the gene products of this region. The *hin* gene encodes an invertase that induces an inversion of the H segment, which includes the *hin* gene itself and the promoter for the *fljB* gene. The *fljB* gene forms an operon with *fljA*, which encodes a negative regulator of *fliC* expression. When the H segment is in the “on” orientation, both *fljA* and *fljB* are transcribed. In contrast, when the H segment is in the “off” orientation, neither gene is transcribed, resulting in the synthesis of phase 1 flagellin (31).

Some serovars of *S. enterica* are described as having R phase or third phase H antigens in the “other” column of the WKL scheme (1). The genetic basis of the expression of these atypical H antigens is known in a few cases. H: j antigen of *S*. Typhi is a variant of its phase 1 antigen d (32). A 261-nucleotide deletion of *fliC* gene is the genetic basis. H:z_66_ antigen of *S*. Typhi is not encoded by neither genes *fliC* nor *fljB*. The gene of which is on a plasmid (33). The third H antigen d of *S*. Rubislaw is encoded by *flpA* gene on the plasmid (32, 34). In this study, we aimed to elucidate the mechanism of R-phase expression in *S*. Senftenberg.

Results of whole-genome sequencing and phylogenetic analysis of *S*. Senftenberg strains revealed that a subset of ST185 strains possess a distinct chromosomal region containing the *fljA*, *fljB*, and *pinR* genes. A deletion mutant lacking this region did not express the H:z_27_ antigen, suggesting that the *fljB* gene encodes the H:z_27_ antigen. Since FljA is known to act as a negative regulator of phase 1 flagellin expression (31), deletion of the *fljA* gene is presumed to result in the expression of the phase 1 flagellin (H:g,s,t). However, we were unable to induce the H:g,s,t antigen in strains HOLHSC-37-53 and HKLHSC-R3-12 by the standard serotyping method. The *pinR* gene is predicted to encode a serine recombinase that mediates inversion of the *pinR* locus and adjacent sequences (35), including the promoter region of the *fljAB* operon. If PinR function is intact, the H:g,s,t antigen may be inducible. The *fljAB* promoter is likely fixed in the “on” orientation, and the frequency of inversion may be too low to allow detection of the alternate H-antigen phase in these strains. Alternatively, mutations within the *pinR* sequence may have led to a loss of function. Indeed, it has been reported that point mutations introduced into the related serine recombinase gene *hin* reduce the frequency of phase variation, thereby preventing induction of the alternate phase of the H antigen (36). For reference, we have not yet succeeded in obtaining a complemented ST185 strain for this region containing the *fljA*, *fljB*, and *pinR* genes.

It is noteworthy that identical or highly similar sequences are distributed among *Escherichia coli* and *Citrobacter* species other than *S*. Senftenberg. In particular, *E. coli* strain EEC-1470 isolated from a case of bovine mastitis harbors a chromosomal region that exhibits 100% sequence identity with the *fljAB-pinR* locus (Accession number: CP010344.1) (37). This finding may suggest that the region was horizontally transferred from bacteria belonging to the order Enterobacterales. The results of the phylogenetic analysis of *S*. Senftenberg isolates belonging various STs suggested that the *fljAB-pinR* locus was transferred to the ST185 lineage around 1867 and spread among this linage. The antigenic formula 1,3,19:z_27_:– was previously designated as serovar Simsbury, but it was later grouped into serovar Senftenberg because conversion to the antigenic formula 1,3,19:g,[s],t:– could be induced(11). Based on the results of the present study, the antigenic formula 1,3,19:g,[s],t:– was shown to represent *S*. Senftenberg ST185 in which the *fljAB-pinR* region was horizontally transferred. Therefore, our findings support the previous decision to unify the name of serovar Simsbury under the serovar Senftenberg.

On the other hand, our phylogenetic analysis of *Salmonella* serovars indicated that ST14 is genetically closer to serovars Albany, Tennessee, and Rissen than to ST185, while ST185 shows a closer relationship to serovar Kentucky than to ST14. As pointed out by Achtman et al., this supports the notion that serotyping does not necessarily reflect evolutionary groupings (38). Nevertheless, since serotyping has long been widely used and provides high discriminatory power, it is considered that the combined use of serotyping and MLST will enable *Salmonella* typing that also reflects the disease potential.

## SUPPLEMENTAL MATERIAL

Supplemental tables. Tables S1 to S6

## DATA AVAILABILITY

The accession information for the strains sequenced in this study is provided in Table S6.

## ACKNOWLEDGEMENTS

This research was supported, in part by the Japan Agency for Medical Research and Development under Grant Number JP24fk0108636.

